# Comparative Analysis of Stain-Free Fourier Ptychographic Microscopy and Traditional Histopathological Light Microscopy in Renal Membranous Nephropathy

**DOI:** 10.1101/2024.09.12.612640

**Authors:** Marika Valentino, Vittorio Bianco, Gioacchino D’Ambrosio, Marco Paulli, Giovanni Smaldone, Valentina Brancato, Lisa Miccio, Marco Salvatore, Marcello Gambacorta, Pietro Ferraro

## Abstract

**Background:** Histology remains a cornerstone in the diagnosis and prognosis of renal diseases, with histopathological analysis of kidney tissue being crucial for understanding renal pathophysiology. The availability of multiple stained sections is essential for conducting a comprehensive histopathological analysis and achieving an accurate diagnosis. Recently, Fourier Ptychographic Microscopy (FPM) earned a spot among the most promising microscopy techniques. The ability to provide high-resolution, quantitative phase-contrast images over a wide area, particularly in a stain-free mode, makes FPM highly appealing to experts in histopathology. Since renal pathologies are characterized by subtle morphological changes encoded in tissue slides, phase maps obtained using FPM are well-suited for providing detailed, high-contrast images of tissue structures. Thus, FPM provides a quantitative imaging tool that can be descriptive of the sample and/or expressive of the disease.

**Methods:** In this study, we explore FPM capability to image pathological kidney tissue, enabling pathologists to select regions of interest within the intricate architecture of renal tissue and zoom in to observe minute submicron structures, ranging from overall tissue organization and glomeruli distribution to individual cell membranes. Attention is focused on membranous glomerulonephritis since it is a nephropathy highly dependent on histological examination.

**Results:** The comparative analysis between FPM and traditional light microscopy showed a difference in thickness of glomerular basal membranes between healthy kidney tissues and those affected by membranous glomerulonephritis (MG). Moreover, the results reported in our investigation revealed better glomerular membranes contrast in FPM images with respect to the H&E-stained images.

**Conclusions:** Our study shows the broad potential of FPM in characterizing hallmarks of MG disease even in stain-free tissue slides.

## 1 Introduction

The histopathological examination of renal tissue biopsy by light microscopy (LM) corroborated by histochemical stains is mandatory to make a diagnosis in kidney disease. The anatomic structure of kidneys, particularly the glomerular complex, has unique morphological features responsible for its physiological functions. Several kidney diseases result in disrupting the anatomy balance and leading to morphological alterations and functional impairment. Thus, modifications of kidney tissue architecture and morphological features may provide diagnostic hallmarks for specific renal disorders.

Kidneys contain numerous functional units called nephrons, each consisting of glomeruli as blood filters, renal tubules as ultra-filters for urine production, and interstitium as support for renal parenchyma and vessels as oxygen and nutrients carriers^1,2^.

The core of renal functioning is represented by the glomerulus. The Bowman’s capsule wraps the glomerulus, which, in turn, encapsulates the capillaries in a sphere delimited by a space, i.e. the Bowman’s space. Epithelial cells and podocytes cover the glomerular capillaries^3^. Endothelial cells maintain the basement membrane of capillaries, which allows filtering water and other substances.

Variations of glomerular structure may include *hypercellularity*, i.e. an increase in glomerular cells, resulting in proliferative glomerulonephritis; *increased extracellular matrix*, i.e. an increase in mesangial or basal membrane material, resulting in basement membrane thickening; *sclerosis*, i.e. a strangling increase in extracellular matrix^1-3^. Some of these morphological alterations may have immunological origin (i.e. induced by deposition of immuno-complex in the capillary wall), or non-immunological origin, i.e. induced by many etiological factors (metabolic and vascular disorders).

Membranous nephropathy (MN) is the most common cause of nephrotic syndrome in adult age^4^, and the main responsible of renal failure within the spectrum of primary glomerulonephritis. Most MN cases (60-80%) are idiopathic, while secondary cases include autoimmune disease, neoplasia, infection and iatrogenic etiology. The pathogenesis of MN consists of deposition of antigen-antibody complexes between the glomerular basement membrane (GBM) and the podocytes, functionally resulting in alterations of the glomerular filtration. The histological hallmark of MN is the occurrence of global thickening and rigidity of the glomerular capillary wall in the absence of signs of cellular proliferation, caused by the deposition of immune complexes, primarily immunoglobulin G (IgG) and complement components^5^.

MN diagnosis and grading of its severity relies on histological evaluation of renal biopsy that is also crucial for monitoring MN disease progression and therapy response. Possible modifications of histological parameters such as glomerular basement membrane thickness, mesangial sclerosis, and interstitial fibrosis can help in treatment decisions and therapeutic interventions.

On such bases, the kidney biopsy seems a suitable candidate for WSI (whole slide imaging) digital pathology application^6,7^. WSI refers to the process of digitalization of “whole” histological slides to create high-resolution digital images that can be viewed, analyzed, and managed by using computer technology. WSI offers numerous advantages, such as accessibility, workflow efficiency, quantitative analysis capabilities, telepathology services, and research applications^6,8^. WSI can facilitate quantitative analysis of histological features through algorithms of image analysis and software tools. Pathologists can perform morphometric measurements, quantify biomarker expression levels, and analyze spatial patterns within tissue specimens to give information about disease pathogenesis, prognosis, and treatment response.

In case of kidney biopsy, Hematoxylin and Eosin (H&E) is one of the most used dyes for renal diseases and identifies structural changes and abnormalities in the glomerular components^9^. Periodic Acid Schiff stain (PAS) marks the glomerulus structures, i.e. basement membrane of the capillary walls^10^. Trichrome – Masson is used to stain the connective tissue and, in case of MN, often reveals subepithelial fuchsinophilic deposits^11^. PAS and Jones methenamine silver (JMS) detect subepithelial spikes formation and/or internal vacuolization of the GBM. Immunofluorescence study can detect deposition of immunoglobulins and complement fractions in the glomerular walls and/or mesangial region^12^.

The typical immunofluorescence feature of MN is the presence of granular subepithelial immune deposits involving the glomerular capillary walls in a diffuse and global pattern. Subepithelial deposits contain IgG in all cases, and complement (C3) in about 80% of cases^3-15^.

However, histological staining procedures may be sometimes associated with morphological tissue changes, including a possible damage to native structures. The quality of the staining also depends on the proper technical procedure employed for dye application, then is often different in the various labs, resulting in limitations in intra- and inter-observer reproducibility^16,17^.

Furthermore, the *conditio sine qua non* for a diagnostic outcome of histological examination is the availability of adequate bioptic material that usually consists of few millimeters of tissue sample. The limited amount of bioptic material coupled with the need of a congruous number of sections for both routinary histology and histochemical stains may often result in a progressive sample exhaustion.

Despite the above reported limitations, a current combination of LM, “in situ” immunofluorescence analyses, and in special cases, electron microscopy, still represents the gold standard in routinary kidney diagnostic pathology^18^.

Among the conventional methodologies^19^, LM has been the first microscopy method to furnish renal structure, alteration, and diseases discrimination^20^. Fluorescence microscopy (FM) provides both detailed architectures and functions of kidney tissue, especially used in glomerulonephritis diagnosis^19,21^. Confocal microscopy (CM) has been used as the first in vivo renal analysis, especially for renal tubular functions^22^. Scanning electron microscopy (SEM) generates high-resolution images that help in observing the intricate renal structures and the ultrastructural alterations^23-26^. Multiphoton microscopy (MPM) permits to study in real-time renal structures and functions, such as glomerular filtration and tubular flow^27^. Cryo-electron microscopy (CEM) provides high-resolution images of renal biomolecules and protein mutations^28^. Transmission electron microscopy (TEM) allows visualizing glomerular and endothelial cells^29^. Notably, most of the above listed microscopy techniques share the need of using staining procedures to observe the inner structures of renal morphology.

In recent decades, stain-free microscopy methodologies have been developed and have rapidly grown, being now well-assessed and mature enough for clinical applications. For instance, Quantitative Phase Imaging (QPI) denotes a wide plethora of microscopy methods for investigating non-invasively and in stain-free mode transparent specimens^30-32^. In QPI, the sample-light interaction depends on the intrinsic properties of the specimens, such as their optical thickness and refractive index^30-32^. The refractive index of biological samples varies according to cell and tissue structures, being linkable to their optical density, and its contrast with respect to the surrounding medium can cause a phase delay within the crossing light beam. Hence, the phase shift due to the presence of the biological object generates the contrast in the QPI images. Such contrast is endogenous, since the specimen is stain-free. The phase-contrast information is linked to quantitative biophysical parameters of the sample, e.g. biovolume or dry mass^33-36^.

Here we use Fourier Ptychographic Microscopy (FPM) to investigate renal diseases^37-40^. FPM allows obtaining high-resolution phase-contrast images over a wide Field of View (FoV), which bypasses the conventional microscopy limits and turns out very useful for tissue analysis^41^.

FPM is based on the synthetic numerical aperture (NA) principle^42^, i.e. a synthesis of multiple NAs, obtained by switching on LEDs of an illumination source array. The LEDs are positioned to generate angled beams of light that probe the sample from various directions in order to transfer in the limited aperture of the microscope objective a wide range of its spatial frequencies; the synthesis of the collected images leverages super-resolution. Typically, wide observation areas are obtained by using a microscope objective with low magnification. The quantitative phase-contrast map is attained by applying phase retrieval algorithms, which reconstruct the sample complex amplitude^37-40^. FPM phase contrast images allowed several studies in biological field. For instance, FPM succeeded in counting white blood cells in blood smear^43^, detecting tumour cells^44^, analysing cells in parallel imaging^45^, providing high-throughput cytometry^46^, observing fibroblast behaviour on wrinkled substrates^47^, studying NIH cells in presence of nanographene particles as multi-scale image^48^, characterizing healthy renal tissues^49^, classifying benign and malign breast cancer by applying fractal geometry on FPM phase images from stain-free paraffined slides^50^. Moreover, FPM promotes the use of artificial intelligence (AI) as few acquisitions produce large datasets for training classifiers^50^ and generative neural networks^51^.

The aim of our research is to test the FPM stain-free method in a series of MN kidney biopsies in order to assess its feasibility and putative diagnostic usefulness in detecting morphological changes in comparison to the traditional histology routine.

## 2 Materials and Methods

### 2.1 Kidney biopsies

Renal biopsies of 9 patients suffering from MN were processed routinely with Logos Milestone microwave processor, cut at 4 µm thickness, de-paraffined and mounted with Sakura Tisse Tek Prisma Plus for the stain-free FPM acquisitions. The same tissue slides were stained in H&E with Sakura Tisse Tek Prisma Plus. All stained slides were scanned with Philips UFS scanner at real magnification of 40x.

The MN diagnosis was made by three expert nephropathologists (MG, GDA, MP) using the Ehrenreich and Chung MN classification^52-54^. In MN, the subepithelial immune deposits progressively accumulate between the glomerular basement membrane and the podocytes, being subsequently incorporated through a series of stages into the GBM. These functional changes correspond to a series of specific histological features detectable by LM, which coupled with immunofluorescence findings can be graded into four stages^52-54^. Stage 1 is defined by the presence of small subepithelial deposits that can be detected exclusively by SEM. In stage 2, the pathological changes are fully developed: glomerular capillary walls are globally thickened, with spike formation separating subepithelial deposits. In this stage the subepithelial immune deposits are electron-dense on SEM images. In stage 3 and 4, the glomerular capillary walls exhibit progressive thickening, due to the incorporation of the deposits into the GBM, resulting in narrowing of the capillary lumens. At these stages the subepithelial immune deposits are electron-lucent on SEM images.

### 2.2 Fourier Ptychographic Microscopy

In FPM, the obtainable lateral resolution depends on the illumination pattern and the NA of the microscope objective^37-42^. All the single LED contributions are synthesized in the Fourier domain, as shown in Fig. 1. Each LED light source allow to record a single image conveying a specific contribution of the sample spatial spectrum. The central LEDs directly illuminate the sample using bright-field illumination (Fig. 1B and 1E). In contrast, each external LED illuminates the sample in ‘dark-field mode’ (Fig. 1C), allowing the acquisition of higher spatial frequencies of the specimen, which fall within the numerical aperture of the microscope objective. As is well known, capturing higher spatial frequencies allows for the reconstruction of fine, high-resolution details using the concept of synthetic aperture retrieval^37^.

**Figure 1.**
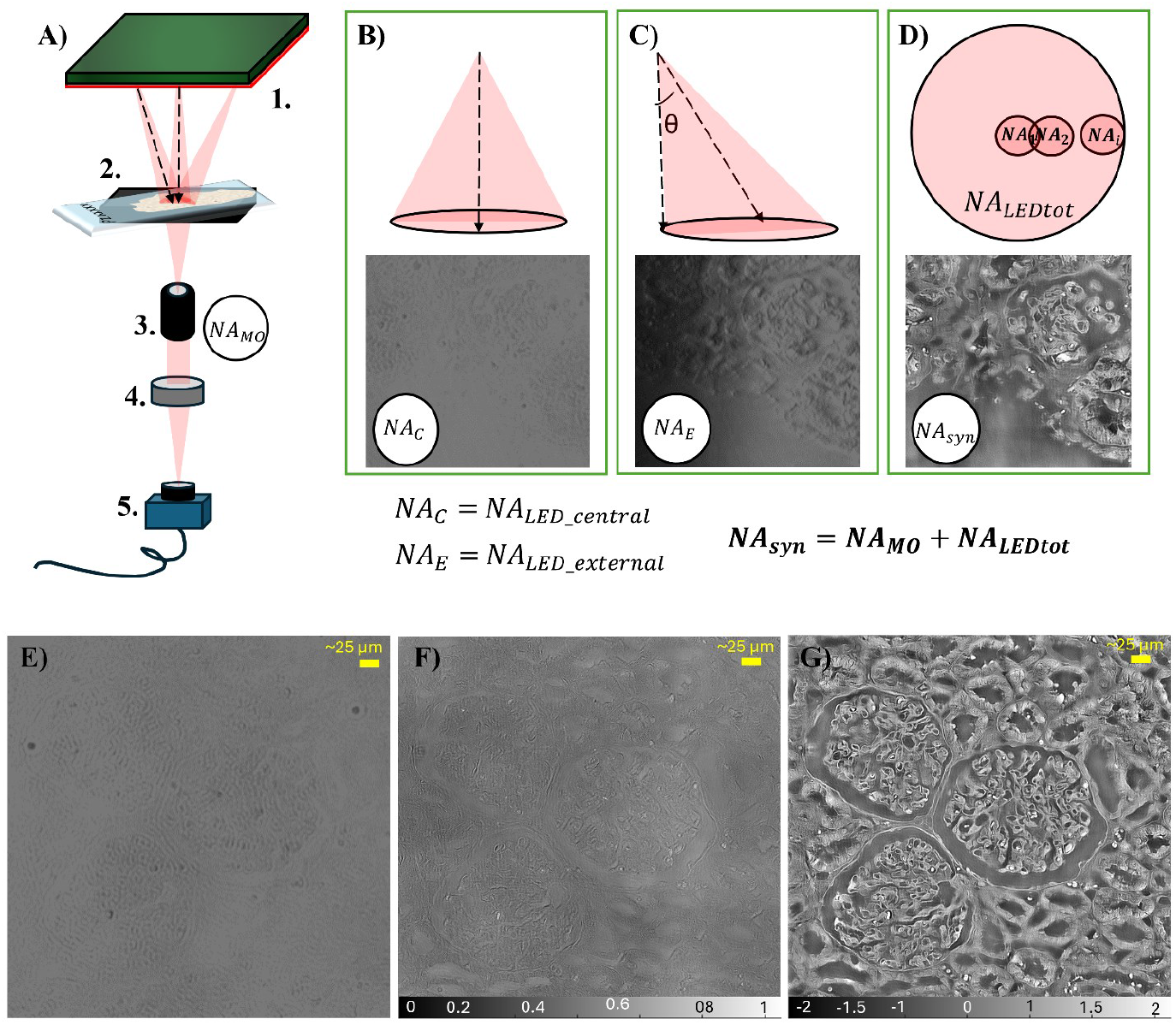
FPM system: A) sketch of the developed FPM setup, where 1. refers to LED illumination array, 2. to the sample holder, 3. to the microscope objective, 4. to the tube lens and 5. to the CCD camera. B) Bright-field image acquired by illuminating the central LED. Dark-field image acquired by a lateral LED illumination angle. D) FPM reconstructed phase-contrast image at high-resolution obtained by synthetizing all LED contributions. E) Example of low-resolution bright-field image of unhealthy glomeruli. F) High-resolution amplitude image of stain-free unhealthy glomeruli. G) High-resolution phase-contrast map of stain-free unhealthy glomeruli.

The synthetic NA radius is determined by the most external LED (Fig. 1 D)). Thus, the synthetic NA is determined as:

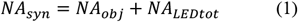

where *NA*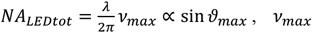 is the maximal spatial frequency that corresponds to the illumination beam with the largest angle, 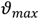, andλis the light wavelength of the FPM system. The lateral resolution of the FPM imaging system improves accordingly:

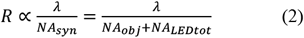

In FPM the captured images are intensity images. Thus, phase retrieval algorithms are needed to obtain the complex object field^37-41^. The complex field of the i-th LED can be simply expressed as 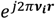 where ***r*** represents the spatial coordinates of the optical system (*x*_*i*_, *y*_*i*_), ***v***_*i*_ the spatial frequency vector and *i* any probing LED of the N-dimensional illumination array. Each LED beam is modulated by the presence of the sample, i.e. the unknown complex object field o(r), and undergoes a phase shift caused by the intrinsic properties of a phase object. Hence, the complex field transmitted by the object assumes the following notation:

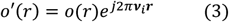

The spatial frequency vector can be expressed in terms of the illumination angle ϑ_*i*_ with respect to the optical axis and the wavelength λ of the FPM system. Thus, in case of tilted illumination, Fig. 1 C), the spatial frequency vector can be written as:

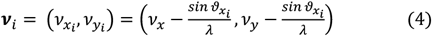

Each image captured by the camera is a low-resolution intensity image, obtained by multiplying the Fourier Transform (FT) of the transmitted complex field, 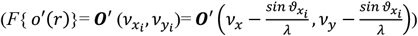 and the transfer function of the optical system (H(*ν*_*x*_, *ν*_*y*_)), i.e. the FT of the impulsive response of the microscope objective (h(r)). The low-resolution (LR) intensity of the tissue sample for each LED is represented by:

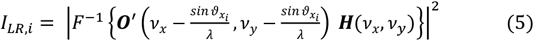

where *F*^−1^ is the inverse Fourier Transform.

As phase retrieval process, here, we use the EPRY (Embedded Pupil function Recovery) algorithm to reconstruct FPM phase contrast images at high lateral resolution^47,55^, i.e. a simultaneous recovery process of Fourier spectra of both object and pupil function. Starting from an initial guess of the high-resolution image (e.g. the i-th LR image, 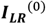), an iterative process of updating the high-resolution image guess is used, where all the LR images are embedded one-by-one. Of course, the algorithm stops when a convergence metric is satisfied. Finally, the high-resolution complex object field (*O*) has the following form:

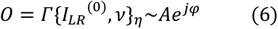

where Γ{…} is the functional operator that resumes the FPM reconstruction process after η iterations (needed for the convergence criterion); *A* is the high-resolution amplitude (Fig. 1 F)) and φ the high-resolution phase-contrast of the observed object (Fig. 1 G)).

### 2.3 FPM experimental setup

Kidney tissue images have been acquired by a custom designed FPM microscope, as shown in Fig. 1 A). A 32×32 RGB LEDs array is used as illumination source. The LEDs array emits a spatially coherent quasi-monochromatic light wave (bandwidth of ∼20 nm). The LEDs are positioned 4 mm aside and 4.67 cm far from the sample stage. The tissue slide is sequentially illuminated by 177 LEDs with red wavelength (632 nm), which follow a circular configuration (with a radius of 60 mm) around the center of the LED matrix. The LED illumination process is controlled by Arduino board and Matlab® scripts. After passing through the object, the diffracted light is filtered by a 4x plan achromatic objective lens (Plan N, 0.1 NA, Olympus) followed by a 400 mm tube lens. The modulated beam reaches a charge-coupled device (CCD) camera (Photometrics Evolve 512), with 4.54 µm pixel pitch and 12-bit quantization. Sample magnification is 4.29x. The low-resolution images size 1460×1940 square pixels.

To promote the plane wave assumption and to fasten the computational time, the LR images are divided into patches^37^. Here, FPM reconstruction process is applied on 100×100 patches of the LR image (i.e. 266 patches in total). The retrieved high-resolution patch has an increased dimension of 500×500 pixels. After unwrapping the phase patches^56^, the entire FoV of the high-resolution image is obtained by stitching all together the 266 patches with an alpha blending algorithm^57^. The final FPM image has a size of 7000×9500 pixels. According to Eq. 1, *NA*_s*yn*_ = 0.1 + 0.5 = 0.6 for our FPM system, thus the lateral resolution reaches 0.51 µm over an area of ∼3 mm^2^ (for further technical details on FPM please refer to Suppl. Mat. of ref. [47]).

## 3 Results

Our FPM imaging system was tested on renal tissue slides from a series of patients affected by MN at different disease stages. Each FPM acquisition was compared to the corresponding LM histologic picture from H&E-stained slide. In Fig. 2, we report an example of LM image (H&E-stained) from a slide of a stage 2 MN and the FPM phase image counterpart. The LM image show a diffuse and regular thickening and rigidity of the glomerular capillary walls (Fig. 2 A-B)); the corresponding FPM high-resolution phase image of the stain-free renal tissue shows higher phase-contrast values in the capillary walls, highlighting the thickening of the GBM (Fig. 2 C-D)). The signal received in the capillary walls is comparable with the traditional histochemical staining. In particular, the pseudo 3D image plot highlights a uniform elevated signal along the capillary wall, consistent with the so-called “railroad track-like thickening” (Fig. 2 E)).

**Figure 2.**
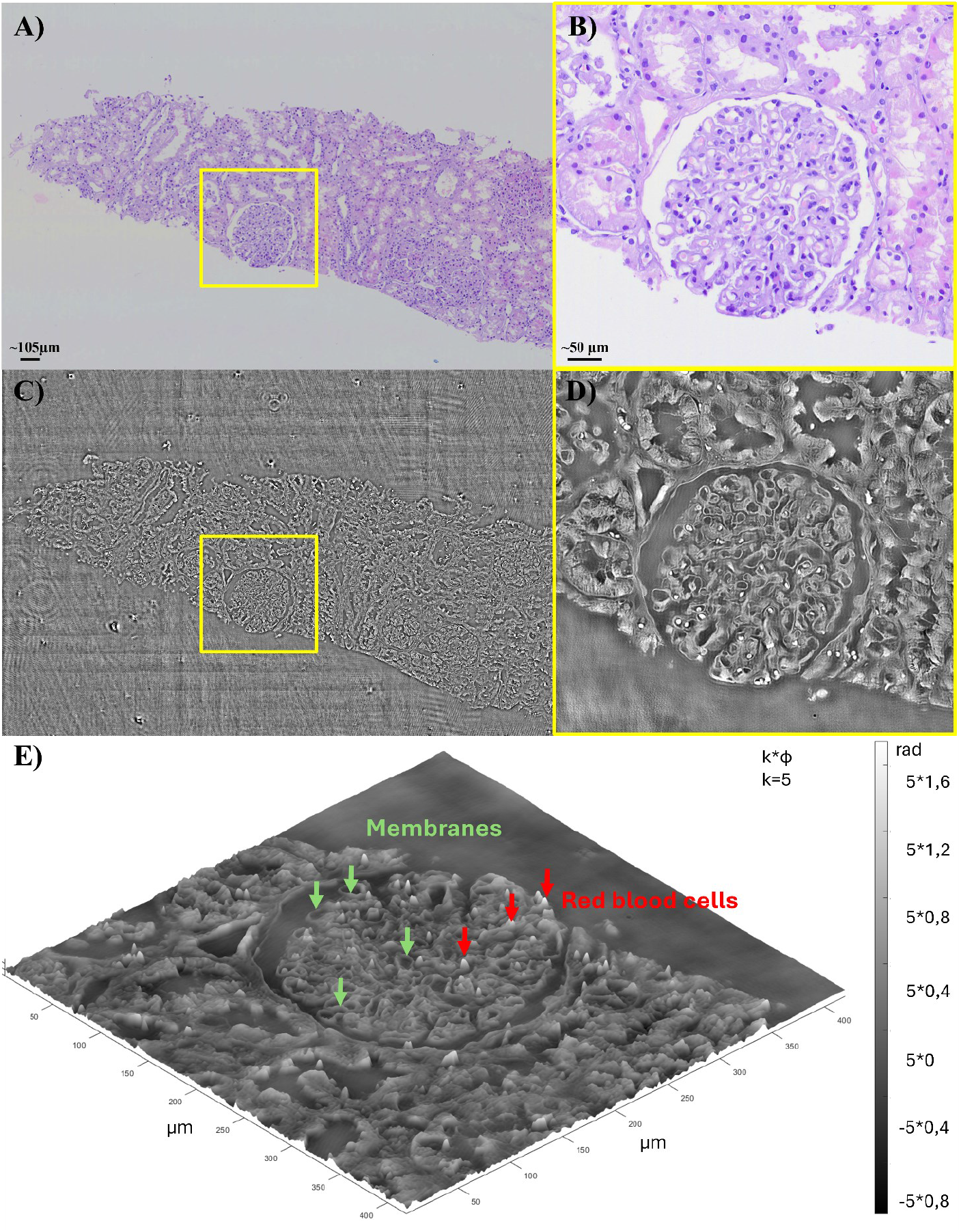
LM vs FPM for MN characterization: A) LM image of H&E-stained renal tissue slide from a case of MN; B) Zoom of a glomerulus with morphological changes of MN (stage 2); C) FPM phase-contrast image of the same area of the unstained renal tissue; Zoom of the same glomerulus of B), FPM image. The maps in C) and D) are shown normalized to the range [0,1]. The phase values after unwrapping are in a range of [-3, 3.5] radians for the entire phase map (C), and [-1.5, 2] radians for the zoomed glomerulus (D). Pseudo 3D image of the FPM phase map, obtained by multiplying the unwrapped phase map by a constant (k=5).

FPM phase image shows a decrease of the signal along the capillary wall, for the sample corresponding to advanced stages of disease. We believe that such lower phase-contrast values, are due to the morphological changes of deposits (yellow arrow Fig. 3 A-B). Several stages of the disease may occur simultaneously in different glomeruli in the same biopsy or different segments of the same glomeruli. For example, in Fig. 3 E-F), the glomerulus shows massive thickening of the capillary walls with initial mesangial sclerosis on the vascular pole (yellow arrow), while the glomerular tip shows less degree of damage (red arrow). The FPM counterpart shows a higher phase-contrast signal on the glomerular tip (red arrow) than in the vascular pole, probably reflecting the different timing of incorporation of the deposits and the different electron-density. Expert pathologists of our team judged these preliminary findings in FPM images very promising: putative advantages include the preservation of tissue sample, the absence of morphological alteration related with staining process, and the availability of morphologic parameters suitable for digital analysis, thus favoring inter-observer standardization.

**Figure 3.**
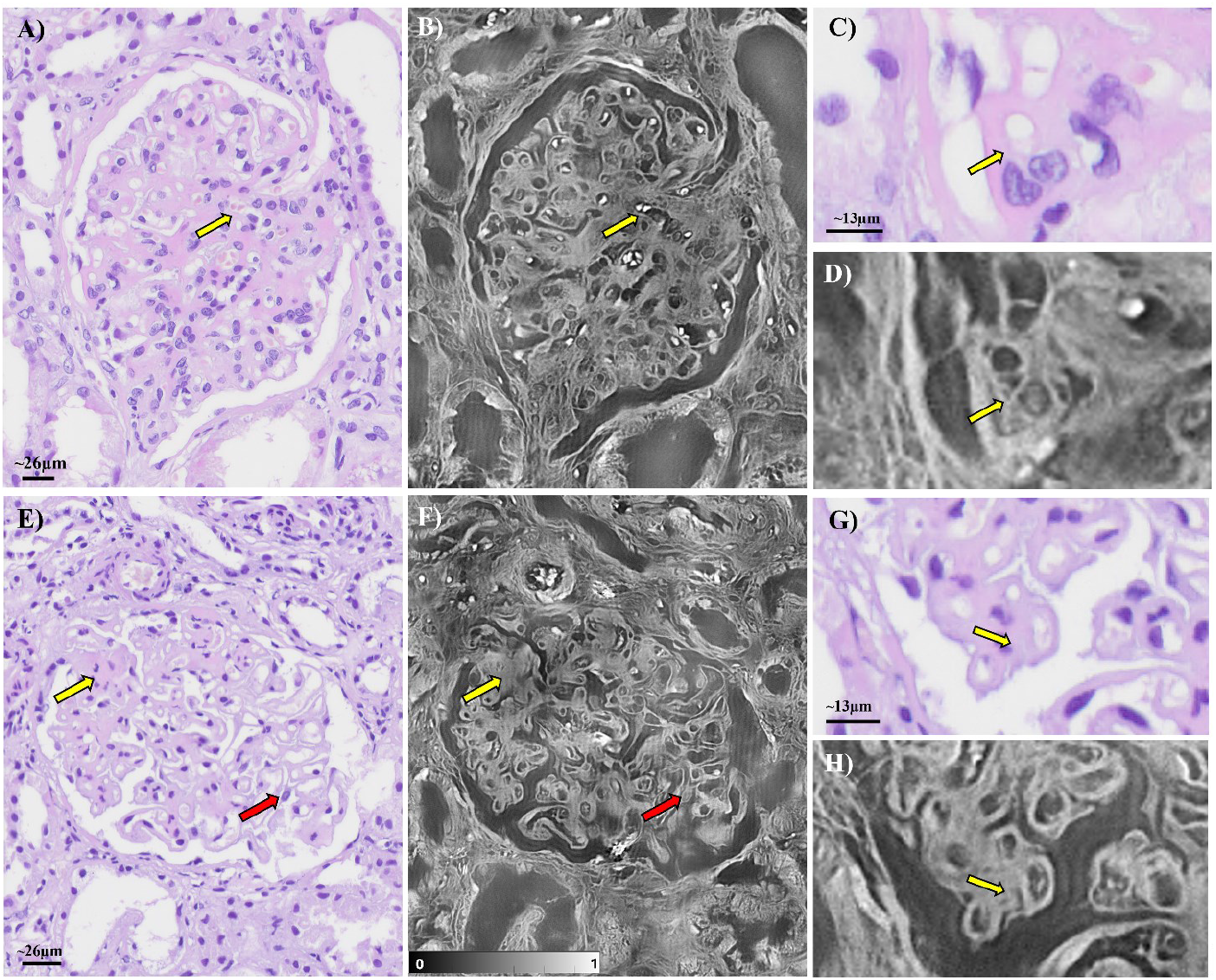
Different stages of the MN pathology: A) LM image, H&E-stained, of a stage 3 MN glomerulus with marked thickening of the capillary walls, narrowing of the capillary lumens and initial mesangial sclerosis; B) FPM phase contrast image of the unstained counterpart. The yellow arrow indicates an area with lower phase contrast that is probably due to the morphological changes of deposits. C-D) Zoomed area of A-B) in which glomerular basal membranes are highly resolved in the FPM phase map. E) LM image, H&E-stained, of two stage variations in the same MN glomerulus: stage 3 in the area indicated by the yellow arrow where there is a mesangial sclerosis on the vascular pole, while stage 2 indicated by the red arrow; F) FPM phase contrast image of the unstained counterpart, where the yellow arrow indicates the 3 stage MN area that has lower phase contrast probably due to the morphological changes of deposits, while the red arrow shows higher phase-contrast values in the capillary walls. G-H) Zoomed area of E-F) where the glomerular basal membranes show the thickening in the FPM phase map. The maps in B) and F) are shown normalized to the range [0,1]. The phase-contrast values after phase unwrapping are in the range [-2, 2] rad for B) and [-2,3] rad for F).

To assess the quantitative nature of the FPM method, and consequently its ability to discriminate renal pathology, we compared the basal membrane profiles of healthy and unhealthy glomeruli in terms of optical thickness (OPT) contrast estimated as:

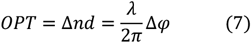

where OPT is the product between the refractive index (RI) contrast, (*n* − *n*_0_), *n*_0_ being the RI of the medium, and the thickness of the observed sample *d*. This product is directly proportional to phase variations Δφ, along with the wavelength of the optical system λ. In Fig. 4 A) the MN glomerulus clearly shows the thickening of the basal membranes. Zooming in an area of the MN glomerulus, we traced a line (light blue) between the basal membranes (Fig. 4 B)) to obtain the OPT profile (Fig. 4 C)). Looking at the plot in Fig. 4 C), the OPT values are reported on the y-axis and show the contrast of optical thickness (along the optical axis) of the glomerular membranes, while the x-axis reports the physical lateral thickness of the glomerular membranes in µm scale. The lateral thicknesses of the considered basal membranes are marked by different colored lines as in the corresponding zoomed image (Fig. 4 B)). We evaluated the maximum and minimum peaks of the OPT distribution as reported in the profile plot in Fig. 4 C). We averaged the maximum peaks and the minimum peaks to estimate the distance between them, i.e. 0.12 µm. Tracing a line (green) on healthy basal membranes (Figs. 4 D-E)), the OPT profile in Fig. 4 F) shows that the healthy basal membranes are significantly thinner than the MN ones. We applied the same analysis to the healthy sample.

**Figure 4.**
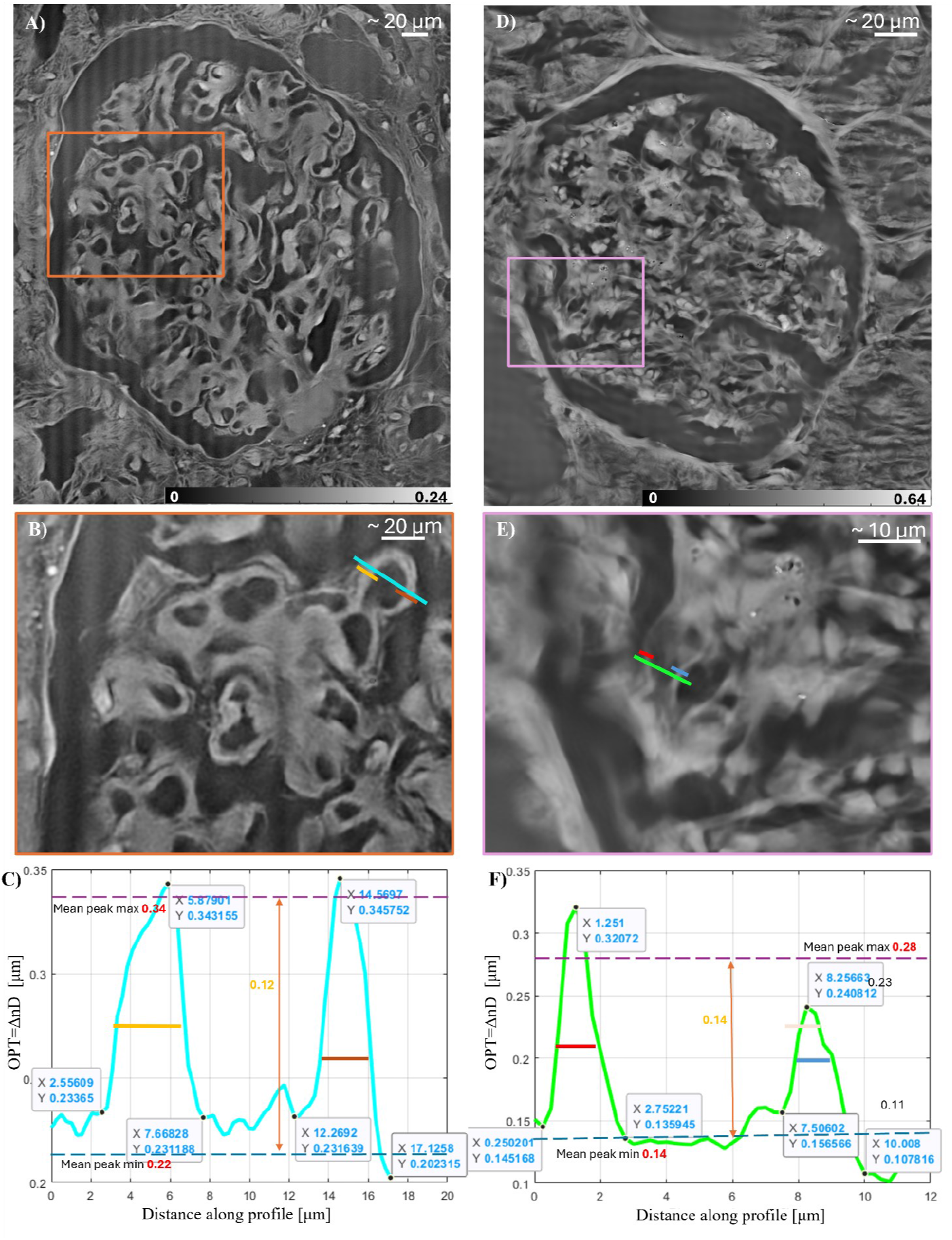
Basal membranes profiles: A) FPM OPT map of MN glomerulus; B) Zoom-in detail showing the basal membranes for the MN case. The coloured line indicates the analysed OPT profile; C) OPT profile of MN basal membranes. D) Healthy glomerulus in FPM OPT map; E) Zoom-in detail of healthy basal membranes. The coloured line indicates the analysed OPT profile; F) OPT profile of healthy basal membranes.

The distance between the mean of the minimum peaks and the mean of the maximum peaks in the healthy basal membranes is 0.14 μm. Furthermore, comparing the values of the lateral thicknesses, the MN basal membranes have a thickness more than 4 μm, while the healthy basal membranes have a thickness less than 2 μm. In Fig. 5, we report more examples of OPT profiles of glomerular basal membranes at different stages of MN and at healthy conditions obtained from OPT maps. Figs. 5 A-D) show the OPT maps of the internal part of glomeruli, whose basal membranes manifest an increase in thickening along with MN stages (from left to right). The colored lines cross the basal membranes to estimate the OPT profiles and their lateral thickness. As in Fig. 4, the maximum peaks indicated in the plot identify each membrane crossed by the profile line. The minimum peaks indicate the boundaries marking the start and end of each membrane. The mean of the maximum peak values and the minimum peak values allow estimating an average of the OPT of the basal membranes of each image. Looking at Figs. 5 A1-D1) the membrane lateral thickness increases according to MN stage. The thickness of the membrane in general ranges between 2 µm to 10 μm in MN cases. In Figs. 5 E-H) we report the OPT contrast maps of the healthy cases with the correlated OPT profiles (Figs. 5 E1-H1). Applying the same analysis, the healthy basal membranes clearly have thinner lateral dimensions, i.e. ∼ 2 µm. Therefore, in the reported examples and in several similar cases not shown here for the sake of conciseness, we found that the lateral thickness clearly increases in the MN case. This can be used as a first biomarker for implementing automatic screening programs or to guide nephrologists.

**Figure 5.**
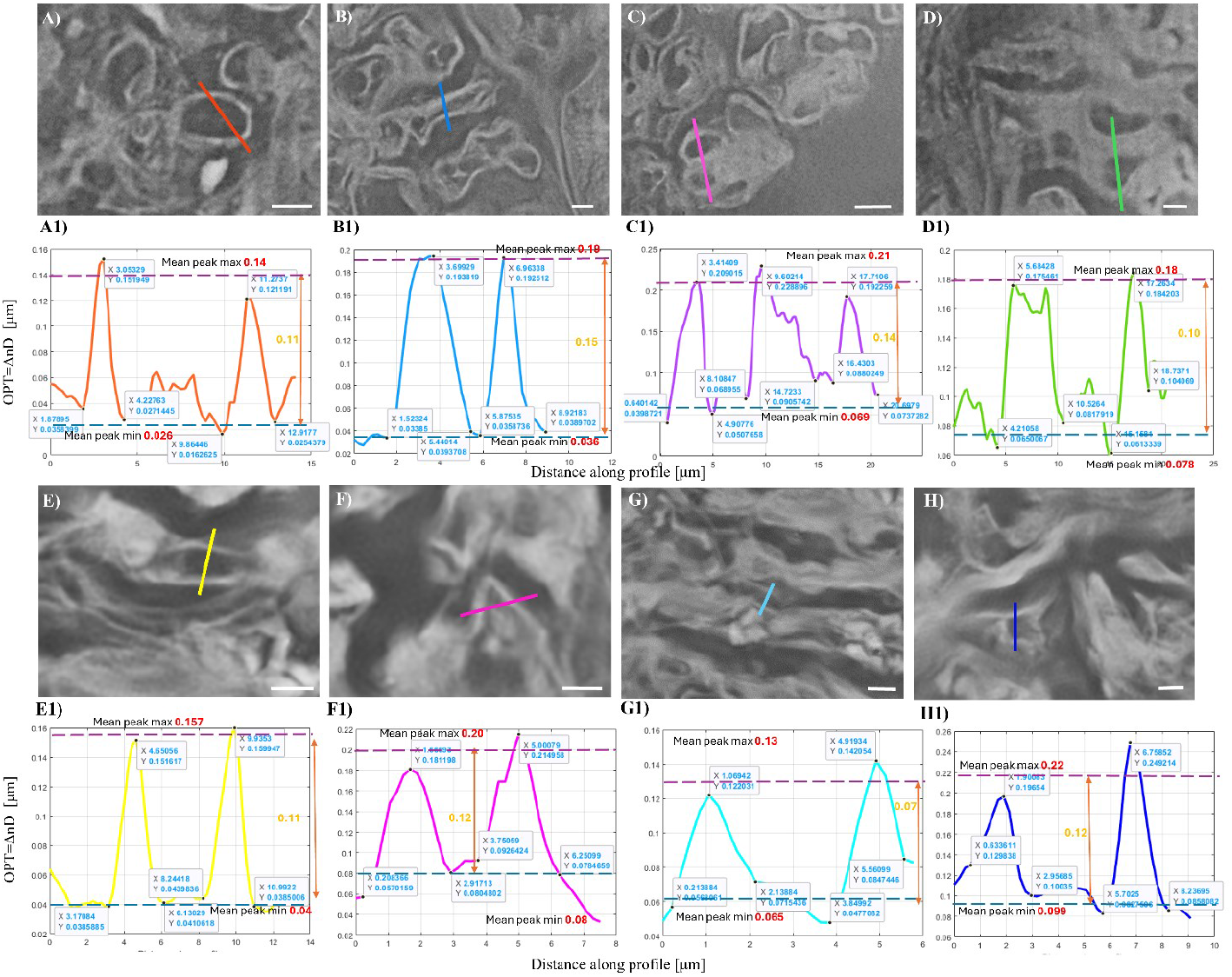
Glomerular membranous profiles in MN cases vs healthy ones: A-D) FPM phase contrast images of internal parts of glomeruli, in particular glomerular basal membranes with MN disease. The thickness increases with the stage of the pathology (from left to right). The lines indicate the profile of OPT. A1-D1) Profile plots of OPT of the previous cases. E-H) FPM phase contrast images of internal parts of glomeruli, in particular glomerular basal membranes in healthy glomeruli. E1-H1) Profile plots of OPT of the previous cases. All the scalebars in A-D) and E-H) are 5 μm sized.

## 4 Discussions

In routinary diagnostic pathology, the analysis of tissue slides is based on observations under conventional optical microscopes. In fact, the final diagnosis is largely based on the pathologist’s expertise corroborated by ancillary techniques including histochemical and immunohistochemical stains, and molecular biology. The recent availability of the digital pathology has provided a further diagnostic aid including machine learning. AI is deeply revolutionizing several fields of biology and medical diagnostics, offering more objective ways of synthesizing large sets of data, improving the performance of standard microscopes, or solving various classification tasks. However, the microscope observations of tissue slides rely on the use of several routinary staining procedures, whose quality is lab-and operator-dependent as previously underlined^16^, resulting in limited inter-labs standardization and possible dissimilar interpretations by the different pathologists as well as AI.

Here, we studied MN in kidney tissue slides in a completely stain-free procedure. Instead of using fluorescence, we used FPM, i.e. a QPI method, to exploit the phase delay as endogenous contrast mechanism. The FPM phase-contrast maps show tiny variations of the local optical density of the various tissue portions, and allows observing the renal tissue at multiple scales, i.e. spanning from its overall organization over a mm^2^ FoV to the smallest traits of the glomeruli with 0,5 *μm* lateral resolution. We have shown that the main features of FPM make it particularly suitable to study nephropathies. We found that the optical resolving power of this computational microscopy method is high enough to study the basal membranes and their alterations in the case of patients with diagnosed MN. The MN tissue slides show thicker membranes than the healthy counterpart, which is very apparent from the analysis of the profiles in Fig. 4 and Fig. 5. This feature is distinctive of the MN and could be used as a quantitative biomarker for setting up automatic classification strategies. These would be based on the stain-free QPI signal rather than the fluorescence readout, making them lab-independent and more reliable and robust. It is worth pointing out that the hardware technology level of many QPI methods and in particular FPM is prone to provide user friendly devices that could be embedded in commercial benchtop microscopes for clinical use in diagnostic laboratories. Likewise, the suite of FPM algorithms is constantly updated to allow non-expert users to work with this type of microscopes, e.g. avoiding complicated calibration steps^58^ and reducing processing times using artificial intelligence for real-time imaging^51^.

In this research, collaboration with expert nephropathologists who evaluated the FPM images enabled an assessment of FPM potential in diagnostics, demonstrating that it can provide image contrast comparable to the stained images typically used in routine diagnosis. The renal morphological changes associated with MN are clearly visible and observable in their inner traits. On such bases, the extraction of FPM features from this type of images might provide an additional tool for corroborating radiomics in clinical diagnosis. For the next future, we expect to extend the FPM analysis to other nephropathies. Further investigations will be planned on larger series of renal biopsies to test the FPM practical diagnostic capability in all different subtypes of nephropathies characterized by abnormal glomerular deposits, e.g. amyloidosis.

## Disclosures

Authors MV, VBi, LM, and PF are employed of the National Research Council (CNR) of Italy. The remaining authors declare that the research was conducted in the absence of any commercial or financial relationships that could be construed as a potential conflict of interest.

## Funding

This work was supported by project BIOLENTI – Sistemi ottici basati su Biolenti in diversi ambiti di ricerca e applicazioni industriali (SOBRI) – of the National Research Council of Italy (Project CUP: B53C22005870005).

## Acknowledgments

The authors gratefully acknowledge Dr. Teresa Rampino, head of Nephrology Unit IRCCS Policlinico San Matteo, who provided the renal biopsies used in this study.

## Author Contributions

MV, VBi, LM, MG, PF, and MS participated in the design of the study. MV performed the experiments. VBr and GS provided H&E-stained LM images. MV analysed data and wrote the original draft. VBi, MG, and PF conceptualized the work. LM, GDA, MP and MG wrote the revised manuscript. VBi, PF and MS supervised the work. All authors read and approved the final manuscript.

## Data Sharing Statement

The data that support the findings of this study are available from the authors upon reasonable request.

## References

1. Bruce M. Koeppen, B.A.S., 2013. Structure and function of the kidneys. Renal Physiology (Fifth Edition), 15–26doi:10.1016/B978-0-323-08691-2.00002-8.

2. Smith, H.W., 1951. The kidney: Structure and function in health and disease. New York: Oxford Univ. Press doi:10.1126/science.114.2969.558.b.

3. Agnes B. Fogo, Arthur H. Cohen, J. Charles Jennette, Jan A. Bruijn, Robert B. Colvin. Fundamentals of renal pathology. © 2006 Springer New York, NY, 1, VI, 222. 10.1007/978-0-387-31127-2.

4. Orth SR, Ritz E. The nephrotic syndrome. N Engl J Med 338:1202–1211, 1998.

5. Schwartz MM. Membranous glomerulonephritis. In: Jennette JC, Olson JL, Schwartz MM, Silva FG, eds. Heptinstall’s Pathology of the Kidney, 5th ed. Philadelphia: Lippincott, 1998:259–308.

6. Evans AJ, Brown RW, Bui MM, Chlipala EA, Lacchetti C, Milner DA, Pantanowitz L, Parwani AV, Reid K, Riben MW, Reuter VE, Stephens L, Stewart RL, Thomas NE. Validating Whole Slide Imaging Systems for Diagnostic Purposes in Pathology. Arch Pathol Lab Med. 2022 Apr 1;146(4):440–450. doi: 10.5858/arpa.2020-0723-CP.

7. Barisoni L, Lafata KJ, Hewitt SM, Madabhushi A, Balis UGJ. Digital pathology and computational image analysis in nephropathology. Nat Rev Nephrol. 2020 Nov;16(11):669–685. doi: 10.1038/s41581-020-0321-6.

8. Kumar N, Gupta R, Gupta S. Whole Slide Imaging (WSI) in Pathology: Current Perspectives and Future Directions. J Digit Imaging. 2020 Aug;33(4):1034–1040. doi: 10.1007/s10278-020-00351-z.

9. Rehan Ahmed Siddiqui, Nurul Kabir, Muhammad Ateeq, Shabana U. Simjee & Muhammad Raza Shah (2016) Characterizing kidney structures in health and diseases using eosin fluorescence from hematoxylin and eosin-stained sections, Journal of Histotechnology, 39:4, 107–115, DOI: 10.1080/01478885.2016.1194608.

10. Matsumoto A, Matsui I, Katsuma Y, Yasuda S, Shimada K, Namba-Hamano T, Sakaguchi Y, Kaimori JY, Takabatake Y, Inoue K, Isaka Y. Quantitative Analyses of Foot Processes, Mitochondria, and Basement Membranes by Structured Illumination Microscopy Using Elastica-Masson- and Periodic-Acid-Schiff-Stained Kidney Sections. Kidney Int Rep. 2021 May 1;6(7):1923–1938. doi: 10.1016/j.ekir.2021.04.021.

11. Cohen AH. Masson’s trichrome stain in the evaluation of renal biopsies. An appraisal. Am J Clin Pathol. 1976 May;65(5):631–43. doi: 10.1093/ajcp/65.5.631.

12. Varis J, Rantala I, Pasternack A. Immunofluorescence of immunoglobulins and complement in kidneys taken at necropsy. J Clin Pathol. 1989 Nov;42(11):1211–4. doi: 10.1136/jcp.42.11.1211.

13. Ehrenreich T, Chung J. Pathology of membranous nephropathy. In: Sommers S, ed. Pathology annual. New York: Appleton-Century-Crofts; 1968.

14. Lai WL, Yeh TH, Chen PM, Chan CK, Chiang WC, Chen YM, Wu KD, Tsai TJ. Membranous nephropathy: a review on the pathogenesis, diagnosis, and treatment. J Formos Med Assoc. 2015 Feb;114(2):102–11.

15. Chung EYM, Wang YM, Keung K, Hu M, McCarthy H, Wong G, Kairaitis L, Bose B, Harris DCH, Alexander SI. Membranous nephropathy: Clearer pathology and mechanisms identify potential strategies for treatment. Front Immunol. 2022 Nov 2;13:1036249.

16. Leica Biosystems. Accessed July 2024. https://www.leicabiosystems.

17. David K. Meyerholz, Amanda P. Beck, Principles and approaches for reproducible scoring of tissue stains in research, Laboratory Investigation, Volume 98, Issue 7, 2018, Pages 844–855, 10.1038/s41374-018-0057-0.

18. D’Agati VD, Mengel M. The rise of renal pathology in nephrology: structure illuminates function. Am J Kidney Dis 2013; 61: 1016–1025.

19. Angelotti ML, Antonelli G, Conte C, Romagnani P. Imaging the kidney: from light to super-resolution microscopy. Nephrol Dial Transplant. 2021 Jan 1;36(1):19–28. doi: 10.1093/ndt/gfz136.

20. Krakower CA. Renal Histopathology: A Light Microscopy Study of Renal Disease. JAMA. 1974;229(1):84–85. doi:10.1001/jama.1974.03230390060037.

21. Buch Archana C, Sood Sonam K, Bamanikar Sunita A, Chandanwale Shirish S, Kumar Harsh, Swapnil Karnik. Role of direct immunofluorescence in the diagnosis of glomerulonephritis. Medical Journal of Dr. D.Y. Patil University. Vol. 8 (4), 452–457, 2015. DOI: 10.4103/0975-2870.160784.

22. Ohno Y, Birn H, Christensen EI. In vivo confocal laser scanning microscopy and micro-puncture in intact rat. Nephron Exp Nephrol 2005; 99: e17–e25.

23. Collan Y, Hirsimäki P, Aho H, Wuorela M, Sundström J, Tertti R, Metsärinne K. Value of electron microscopy in kidney biopsy diagnosis. Ultrastruct Pathol. 2005 Nov-Dec;29(6):461–8. doi: 10.1080/01913120500323381.

24. de Kort H, Moran L, Roufosse C. The role of electron microscopy in renal allograft biopsy evaluation. Curr Opin Organ Transplant. 2015 Jun;20(3):333–42. doi: 10.1097/MOT.0000000000000183.

25. Strong ML, Evers P. The third dimension in renal diagnosis. Scanning electron microscopy of normal and abnormal kidney. S Afr Med J 1979; 55: 174–177.

26. Liapis H. Electron microscopy in kidney research: seeing is believing. Ultrastruct Pathol 2013; 37: 340–345.

27. Hackl MJ, Burford JL, Villanueva K et al. Tracking the fate of glomerular epithelial cells in vivo using serial multiphoton imaging in new mouse models with fluorescent lineage tags. NatMed 2013; 19: 1661–1666.

28. Fernandez-Leiro R, Scheres SH. Unravelling biological macromolecules with cryo-electron microscopy. Nature 2016; 537: 339–346.

29. Hegermann, J., Lünsdorf, H., Ochs, M. et al. Visualization of the glomerular endothelial glycocalyx by electron microscopy using cationic colloidal thorium dioxide. Histochem Cell Biol 145, 41–51 (2016). 10.1007/s00418-015-1378-3.

30. Popescu, G., (2011). Quantitative Phase Imaging of Cells and Tissues. McGraw-Hill Professional: New York, NY, USA.

31. Park, Y., Depeursinge, C. and Popescu, G., (2018). Quantitative phase imaging in biomedicine. Nature Photon 12, 578–589. 10.1038/s41566-018-0253-x.

32. Thang L. Nguyen, Soorya Pradeep, Robert L. Judson-Torres, Jason Reed, Michael A. Teitell and Thomas A. Zangle. Quantitative Phase Imaging: Recent Advances and Expanding Potential in Biomedicine. ACS Nano 2022 16 (8), 11516–11544, DOI: 10.1021/acsnano.1c11507.

33. Zangle, T. A.; Teitell, M. A. Live-Cell Mass Profiling: An Emerging Approach in Quantitative Biophysics. Nat. Methods 2014, 11 (12), 1221–8.

34. Ganoza-Quintana, J.L.; Arce-Diego, J.L.; Fanjul-Vélez, F. Digital Histopathological Discrimination of Label-Free Tumoral Tissues by Artificial Intelligence Phase-Imaging Microscopy. Sensors 2022, 22, 9295. 10.3390/s22239295.

35. Zangle, T. A.; Teitell, M. A.; Reed, J. Live Cell Interferometry Quantifies Dynamics of Biomass Partitioning During Cytokinesis. PLoS One 2014, 9 (12), e115726.

36. B. Rappaz, P. Marquet, E. Cuche, Y. Emery, C. Depeursinge, and P. J. Magistretti, “Measurement of the integral refractive index and dynamic cell morphometry of living cells with digital holographic microscopy,” Opt. Express 13, 9361–9373 (2005).

37. Zheng, G., Horstmeyer, R. & Yang, C. (2013). Wide-field, high-resolution Fourier ptychographic microscopy. Nature Photon 7, 739–745. 10.1038/nphoton.2013.187.

38. Konda Pavan Chandra, Loetgering Lars, Zhou Kevin C., Xu Shiqi, Harvey Andrew R., and Horstmeyer Roarke, (2020). Fourier ptychography: current applications and future promises. Opt. Express 28, 9603–9630. 10.1364/OE.386168.

39. Xu, F.; Wu, Z.; Tan, C.; Liao, Y.; Wang, Z.; Chen, K.; Pan, A. Fourier Ptychographic Microscopy 10 Years on: A Review. Cells 2024, 13, 324. 10.3390/cells13040324.

40. Ou X., Horstmeyer R., Yang C., and Zheng G., (2013). Quantitative phase imaging via Fourier ptychographic microscopy. Opt. Lett., Vol. 38, pp. 4845–4848. doi: 10.1364/OL.38.004845.

41. Shuhe Zhang, Aiye Wang, Jinghao Xu, Tianci Feng, Jinhua Zhou, An Pan. FPM-WSI: Fourier ptychographic whole slide imaging via feature-domain backdiffraction. arXiv:2402.18270, 2024. 10.48550/arXiv.2402.18270.

42. Ou X, Horstmeyer R, Zheng G, Yang C. High numerical aperture Fourier ptychography: principle, implementation and characterization. Opt Express. 2015 Feb 9;23(3):3472–91. doi: 10.1364/OE.23.003472.

43. Chung J, Ou X, Kulkarni RP, Yang C. Counting White Blood Cells from a Blood Smear Using Fourier Ptychographic Microscopy. PLoS One. 2015 Jul 17;10(7):e0133489. doi: 10.1371/journal.pone.0133489.

44. Williams A, Chung J, Ou X, Zheng G, Rawal S, Ao Z, Datar R, Yang C, Cote R. Fourier ptychographic microscopy for filtration-based circulating tumor cell enumeration and analysis. J Biomed Opt. 2014 Jun;19(6):066007. doi: 10.1117/1.JBO.19.6.066007.

45. Chan, A.C.S., Kim, J., Pan, A. et al. Parallel Fourier ptychographic microscopy for high-throughput screening with 96 cameras (96 Eyes). Sci Rep 9, 11114 (2019). 10.1038/s41598-019-47146-z.

46. Chengfei Guo, Shaowei Jiang, Liming Yang, Pengming Song, Azady Pirhanov, Ruihai Wang, Tianbo Wang, Xiaopeng Shao, Qian Wu, Yong Ku Cho, Guoan Zheng, Depth-multiplexed ptychographic microscopy for high-throughput imaging of stacked bio-specimens on a chip, Biosensors and Bioelectronics, Volume 224, 2023, 115049, 10.1016/j.bios.2022.115049.

47. Pirone D., Bianco V., Valentino M., Mugnano M., Pagliarulo V., Memmolo P., Miccio L., Ferraro P., (2022). Fourier ptychographic microscope allows multi-scale monitoring of cells layout onto micropatterned substrates. Optics and Lasers in Engineering, Volume 156, 107103, 10.1016/j.optlaseng.2022.107103.

48. Marika Valentino et al. QPI assay of fibroblasts resilience to adverse effects of nanoGO clusters by multimodal and multiscale microscopy. J. Phys. Photonics 6 015004, 2024. DOI 10.1088/2515-7647/ad1c6b.

49. Valentino M, Bianco V, Miccio L, Memmolo P, Brancato V, Libretti P, Gambacorta M, Salvatore M and Ferraro P (2023) Beyond conventional microscopy: Observing kidney tissues by means of fourier ptychography. Front. Physiol. 14:1120099. doi: 10.3389/fphys.2023.1120099.

50. Vittorio Bianco, Marika Valentino, Daniele Pirone, Lisa Miccio, Pasquale Memmolo, Valentina Brancato, Luigi Coppola, Giovanni Smaldone, Massimiliano D’Aiuto, Gennaro Mossetti, Marco Salvatore, Pietro Ferraro. Classifying breast cancer and fibroadenoma tissue biopsies from paraffined stain-free slides by fractal biomarkers in Fourier Ptychographic Microscopy. Computational and Structural Biotechnology Journal, Vol. 24, 225–236 2024. 10.1016/j.csbj.2024.03.019.

51. Francesco Bardozzo, Pierpaolo Fiore, Marika Valentino, Vittorio Bianco, Pasquale Memmolo, Lisa Miccio, Valentina Brancato, Giovanni Smaldone, Marcello Gambacorta, Marco Salvatore, Pietro Ferraro, Roberto Tagliaferri, Enhanced tissue slide imaging in the complex domain via cross-explainable GAN for Fourier ptychographic microscopy, Computers in Biology and Medicine, Volume 179, 2024, 108861, 10.1016/j.compbiomed.2024.108861.

52. Ehrenreich T, Chung J. Pathology of membranous nephropathy. In: Sommers S, ed. Pathology annual. New York: Appleton-Century-Crofts; 1968.

53. Lai WL, Yeh TH, Chen PM, Chan CK, Chiang WC, Chen YM, Wu KD, Tsai TJ. Membranous nephropathy: a review on the pathogenesis, diagnosis, and treatment. J Formos Med Assoc. 2015 Feb;114(2):102–11.

54. Chung EYM, Wang YM, Keung K, Hu M, McCarthy H, Wong G, Kairaitis L, Bose B, Harris DCH, Alexander SI. Membranous nephropathy: Clearer pathology and mechanisms identify potential strategies for treatment. Front Immunol. 2022 Nov 2; 13:1036249.

55. Xiaoze Ou, Guoan Zheng, and Changhuei Yang, “Embedded pupil function recovery for Fourier ptychographic microscopy,” Opt. Express 22, 4960–4972 (2014).

56. Yuan Guo, Xiaotian Chen, Tao Zhang, Robust phase unwrapping algorithm based on least squares, Optics and Lasers in Engineering, Volume 63, 2014, Pages 25–29, 10.1016/j.optlaseng.2014.06.007.

57. Jiakun Bao, Jingtao Fan, X.H.J.W.L.W., 2017. An effective consistency correction and blending method for camera-array-based microscopy imaging. International Conference on Systems, Signals and Image Processing (IWSSIP).

58. V. Bianco et al., “Deep Learning-Based, Misalignment Resilient, Real-Time Fourier Ptychographic Microscopy Reconstruction of Biological Tissue Slides,” in IEEE Journal of Selected Topics in Quantum Electronics, vol. 28, no. 4: Mach. Learn. in Photon. Commun. and Meas. Syst., pp. 1–10, July-Aug. 2022, Art no. 6800110, doi: 10.1109/JSTQE.2022.3154236.

